# Cell-to-cell variability of dynamic CXCL12-CXCR4 signaling and morphological processes in chemotaxis

**DOI:** 10.1101/2022.05.19.492090

**Authors:** Kenneth K.Y. Ho, Siddhartha Srivastava, Patrick C. Kinnunen, Krishna Garikipati, Gary D. Luker, Kathryn E. Luker

## Abstract

Chemotaxis drives critical processes in cancer metastasis. While commonly studied at the population scale, metastasis arises from small numbers of cells that successfully disseminate, underscoring the need to analyze chemotaxis at single-cell resolution. Here we focus on chemotaxis driven by the CXCL12-CXCR4 pathway, a signaling network that promotes metastasis in more than 20 different human cancers. CXCL12-CXCR4 activates ERK and Akt, kinases known to promote chemotaxis, but how cells couple signaling to chemotaxis remain poorly defined. To address this challenge, we implemented single-cell analysis of MDA-MB-231 breast cancer cells migrating in a chemotaxis device towards chemokine CXCL12. We integrated live, single-cell imaging with advanced computational analysis methods to discover processes defining subsets of cells that move efficiently toward a CXCL12 gradient. We identified dynamic oscillations in ERK and Akt signaling and associated morphological transitions as key determinants of successful chemotaxis. Cells with effective chemotaxis toward CXCL12 exhibit faster and more persistent movement than non-migrating cells, but both cell populations show similar random motion. Migrating cells exhibit higher amplitude fluctuations in ERK and Akt signaling and greater frequencies of generating lateral cell membrane protrusions. Interestingly, computational analysis reveals less correlated network coupling of signaling and morphological changes in migrating cells. These data reveal processing events that enable cells to convert a signaling input to chemotaxis and highlight how cells in a uniform environment produce heterogeneous responses.

## Introduction

Chemokine CXCL12 and its receptor, CXCR4, contribute to organ-specific chemotaxis of circulating tumor cells and subsequent metastasis in breast cancer and multiple other malignancies (1, 2). Crucial functions of CXCL12-CXCR4 signaling in metastatic cancer motivate ongoing efforts to target this signaling pathway with novel therapies (3). Studies of CXCL12-CXCR4 in cancer and compounds targeting this pathway commonly assume that all cells expressing CXCR4 signal and respond to CXCL12. However, our group discovered marked heterogeneity in activation of intracellular signaling and chemotaxis among seemingly identical CXCR4-positive cancer cells treated with CXCL12 (4, 5). Such heterogeneity is not unique to CXCL12-CXCR4 as prior studies report similar responses to other signaling molecules, such as EGF and EGFR (6-9). These responses are not simply due to stochasticity but contain governing rules behind (10, 11). The fact that cells even in two-dimensional culture can express a receptor yet fail to respond has critical implications for development of molecular biomarkers and targeted therapies. The mere presence of a receptor may not correlate with a cancer cell consistently relying on that molecule for oncogenic signaling and functions. Therefore, successfully targeting that receptor might fail to block tumor progression and metastasis. Lack of insights about mechanisms that regulate heterogeneous responses of cancer cells remains a critical barrier as even a single cancer cell that evades therapy could potentially lead to metastatic disease.

CXCL12-CXCR4 signaling activates downstream PI3K/Akt and MAPK/ERK pathways. Both Akt and ERK kinases promote reorganization of the actin cytoskeleton in cells with resultant changes in cell morphology needed for cell migration and chemotaxis (12, 13). This process is driven by feedbacks between actin cytoskeleton and PI3K/MAPK signaling resulting an oscillatory network regulating cell migration. Fluctuation of this oscillatory network occurs rapidly on the timescale of minutes after stimulating cells with CXCL12, while chemotaxis toward a gradient of CXCL12 requires several hours to detect directional movement. Processes cells use to connect signaling, morphology, and chemotaxis across these timescales remain incompletely understood, particularly in the context of how heterogeneity in signaling directly relates to heterogeneous chemotaxis among single cells.

To investigate single-cell heterogeneity in signaling and chemotaxis toward CXCL12, we employed an integrated experimental approach combining live, single-cell imaging and advanced computational analysis methods. We studied to what extend quantitative interconnections of ERK and Akt signaling dynamics and morphological transitions control migratory MDA-MB-231 breast cancer cells. We implemented multiplexed fluorescence reporters with automated live-cell imaging and image processing to quantify dynamic signaling, morphology, and chemotaxis data in >1000 cells over 24 hours at 4-minute time intervals. Next, we used a time-dependent data processing approach to extract features that quantify dynamic processes in the multidimensional data. Then, we devise a physics-based system inference approach to identify data-driven chemotaxis properties. Further, we used dimensionality reduction techniques and cross-correlation analysis to quantify the relationship of signaling, morphology, and chemotaxis. We found differences between migratory and non-migratory cells in their speed and persistence, but not random motion. We also discovered ERK and Akt signal fluctuations and cell morphological transition frequencies both associating with the migratory subpopulation. Lastly, we found a less correlated network connecting ERK and Akt signals and morphological transition in migrating cells. These data provide previously unidentified insights in unique signaling and morphological interconnectedness in migrating cell subpopulation.

## Results

### Heterogeneous and dynamic networks controlling chemotaxis

Revealing potential causes of heterogeneity in dynamic network connecting signaling, morphology, and chemotaxis in cells calls for a large multidimensional dataset. One that includes multiplexed inputs, hundreds of time points, and hundreds of single-cell observations; and handles heterogeneous cell movement and mitotic events over time. We stably expressed multiplexed fluorescent reporters, histone 2B fused to mCherry (H2B-mCherry) and independent Akt and ERK kinase translocation reporters (KTRs), in cells to capture signaling, morphology, and chemotaxis information. H2B-mCherry marks cell nuclei, quantifies cell chemotactic movement (**Fig. 1A-i**), and enables image segmentation. ERK and Akt KTRs enable measurements of respective kinase activities (14, 15) and mark cell shape. The KTRs are designed to translocate reversibly out of the nucleus when the respective kinase is active (**Fig. 1A-ii**). Quantifying fluorescence intensity ratios in the cytoplasm to nucleus provides analog and independent measurements of ERK and Akt kinase activities in single cells. Further, segmenting the cell based on combined ERK and Akt KTR channels and best-fitting with an ellipse provide quantitative measurements of single-cell morphological properties (**Fig. 1A-iii**). Cells expressing multiplexed fluorescent reporters were conditioned to chemotax in the microfluidic device and were imaged over time. Microfluidic device has steady control of CXCL12 chemotactic gradient over time. It also provides favorable environment for long term migration of cells and enables automated live-cell fluorescence imaging (**Fig. S1**). Fluorescence images were processed to generate single-cell multiplexed data using our automated image processing pipeline with identification of mitotic events and morphology assisted tracking.

**Figure 1:**
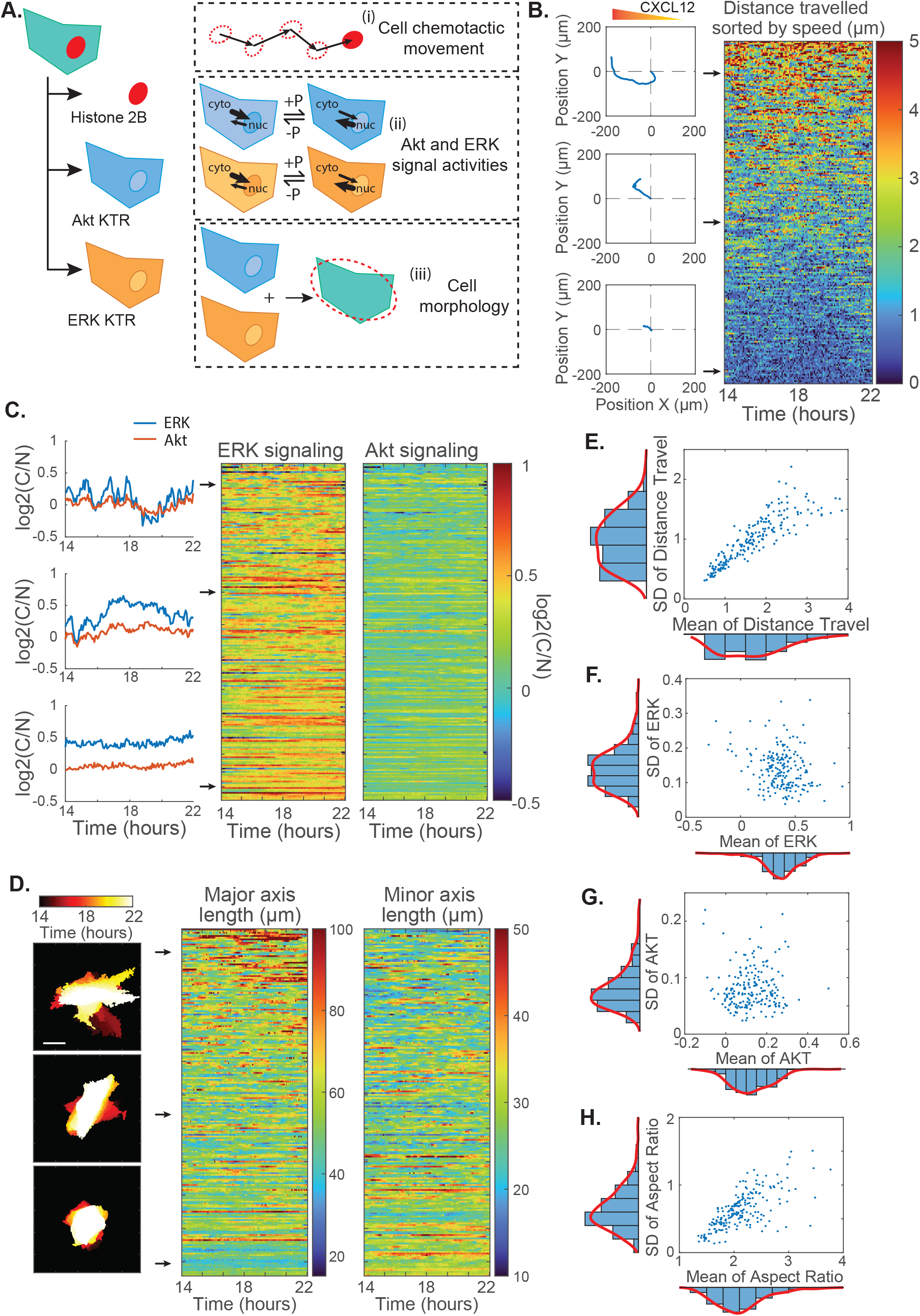
Heterogeneous and dynamic processes in MDA-MB-231 cells chemotaxis under CXCL12 gradient. **A**. Schematic showing the multiplexed fluorescence reporters: histone 2B marker, Akt kinase translocation reporter (KTR), and ERK KTR. These fluorescence reporters were used to quantify cell chemotactic movements (i), Akt and ERK signal activities (ii), and cell morphology (iii). **B**. Pseudocolor plot shows the heterogeneous and dynamic travelling distance across 8 hours of example in the whole 24-hour study. Three example trajectories on the left show cell-to-cell variation in migration behaviors. **C**. Pseudocolor plots show heterogeneous and dynamic fluctuation of ERK and Akt signaling across 8 hours of example data from the whole 24-hour study. Three examples on the left show ERK and Akt signals vary from large fluctuation amplitudes to negligible fluctuations. **D**. Pseudocolor plots show heterogeneous changes in cell shape, as represented in major and minor axis lengths. Three examples on the left show different degree of changes in cell segmented mask centered at nuclear centroid over 8 hours of time. Scale bar = 25µm. **E-H**. Scatter-histogram plots show the relationship and distribution of means and standard deviations of movement (E), ERK (F) and Akt (G) signaling, and morphology (H). A non-linear relationship represents absolute levels and fluctuation amplitudes as important sources of heterogeneity. Each point represents an individual cell in the experiment. (8 hours of data capturing 208 cells taken every 4 minutes are shown as an example out of a total of 24 hours of data with >1000 cells)

After quantifying cell movement, morphology, and ERK and Akt signaling in single cells, we observed heterogeneous and dynamic responsiveness in MDA-MB-231 breast cancer cells expressing CXCR4 receptor under CXCL12 gradient as the only stimulus. Aligned with previous study showing heterogeneous chemotaxis response to CXCL12 (4), single-cell chemotaxis behaviors ranged from migrators to non-migrators based on average travelling speed despite expression of CXCR4 (**Fig. 1B left**). Time-dependent information in cell’s position reveal that distance travelled by individual cell is not uniform but fluctuates over time. For example, individual cells switch between high travelling speed mode and small movement mode repeatedly in 8 hours of time (**Fig. 1B right**). We ask to what extent signaling and morphology changes contribute to heterogeneous and dynamic chemotactic movement. We found that cells also exhibit fluctuations in ERK and Akt signaling, similar to what were reported in previous studies, driving cell behaviors like cell division, death, and migration (12, 13, 16-20). Single-cell signaling responses varied from strong fluctuations of both ERK and Akt to undetectable fluctuations (**Fig. 1C**). Despite constitutively active ERK signaling, MDA-MB-231 cells exhibit greater fluctuations of ERK than Akt, suggesting fluctuation amplitudes may not relate to overall kinase activities. Oscillatory network coupling ERK/Akt signaling and actin cytoskeleton results changes in cell morphology during chemotaxis (21, 22). Quantification of cell shape using aspect ratio of major and minor axis lengths reveals single-cell morphological responses ranged from strong fluctuations of major and minor axis lengths to negligible fluctuations (**Fig. 1D**).

Apart from the highlighted cell-to-cell variations in cell movement, signaling, and morphology fluctuations, initial and average values also show a range of deviations among cells in our single-cell analysis, as reported in past works (5, 7, 23-25). We ask to what extent fluctuation is an independent descriptor of heterogeneity on top of cell-to-cell variability in initial and average values (**Fig. 1E-H**). Lack of correlation between the mean and standard deviation for signaling (correlation coefficient of -0.24 for ERK and 0.01 for Akt) indicates that both absolute level and fluctuation amplitude of ERK and Akt are independent descriptors of heterogeneity among individual cells (**Fig. 1F-G**). Conversely, stronger correlations between the mean and standard deviation in migration and morphology behavior (0.75 for distance travelled and 0.73 for aspect ratio) suggest that fluctuations in migration and morphology behavior are not as important as signaling fluctuations (**Fig. 1E, H**). This suggests the importance of analyzing both dynamics and heterogeneity of cancer cell responsiveness at the same time, especially for ERK and Akt signaling during chemotaxis. Therefore, we seek to investigate to what extent the dynamics of signaling and morphological changes relate to migration heterogeneity.

### Unique traits of migrating cells

Quantified from automated image processing, chemotaxis data are 2-dimensional information for nuclear positions that involve changes in speed and direction over time. Therefore, before we focus on finding unique signaling and morphology determinants for successful chemotaxis, a non-biased method is needed to separate cells into migrating vs non-migrating cells. Previous studies have defined migration features to describe chemotaxis behavior, such as speed, persistence, accuracy, and velocity in x and y directions (21, 23, 26, 27). Speed is the accumulated distance travelled for each cell over time. Persistence is the ratio between displacement and distance and depicts how straight each cell is moving. Accuracy is cosine of an angle between final position and original position of a cell with respect to the axis of chemotactic gradient. Velocity in the x and y direction is the velocity component parallel and perpendicular to the chemotactic gradient respectively. However, since these features are related to each other (23), any of these migration features alone cannot help determine whether individual cell is migratory or not. To characterize heterogeneous and dynamic chemotaxis behaviors, we implemented dimensionality reduction techniques and data-driven computational analysis to separate cells into subpopulations and compare behaviors of migrating and non-migrating cells (**Fig. 2A**). More than 1200 single-cell MDA-MB-231 data from two independent CXCL12 mediated chemotaxis repeats are separated into three subpopulations: 28.5% of cells belong to cluster 1 and migrated towards CXCL12, 35.2% of cells belong to cluster 2 and migrated away from CXCL12 source, and 36.3% of cells are non-migratory (cluster 3) and stayed around their original location (**Fig. 2B-C**). To further quantify the heterogeneous chemotaxis response, we ask what unique traits separate cells from different subpopulations.

**Figure 2:**
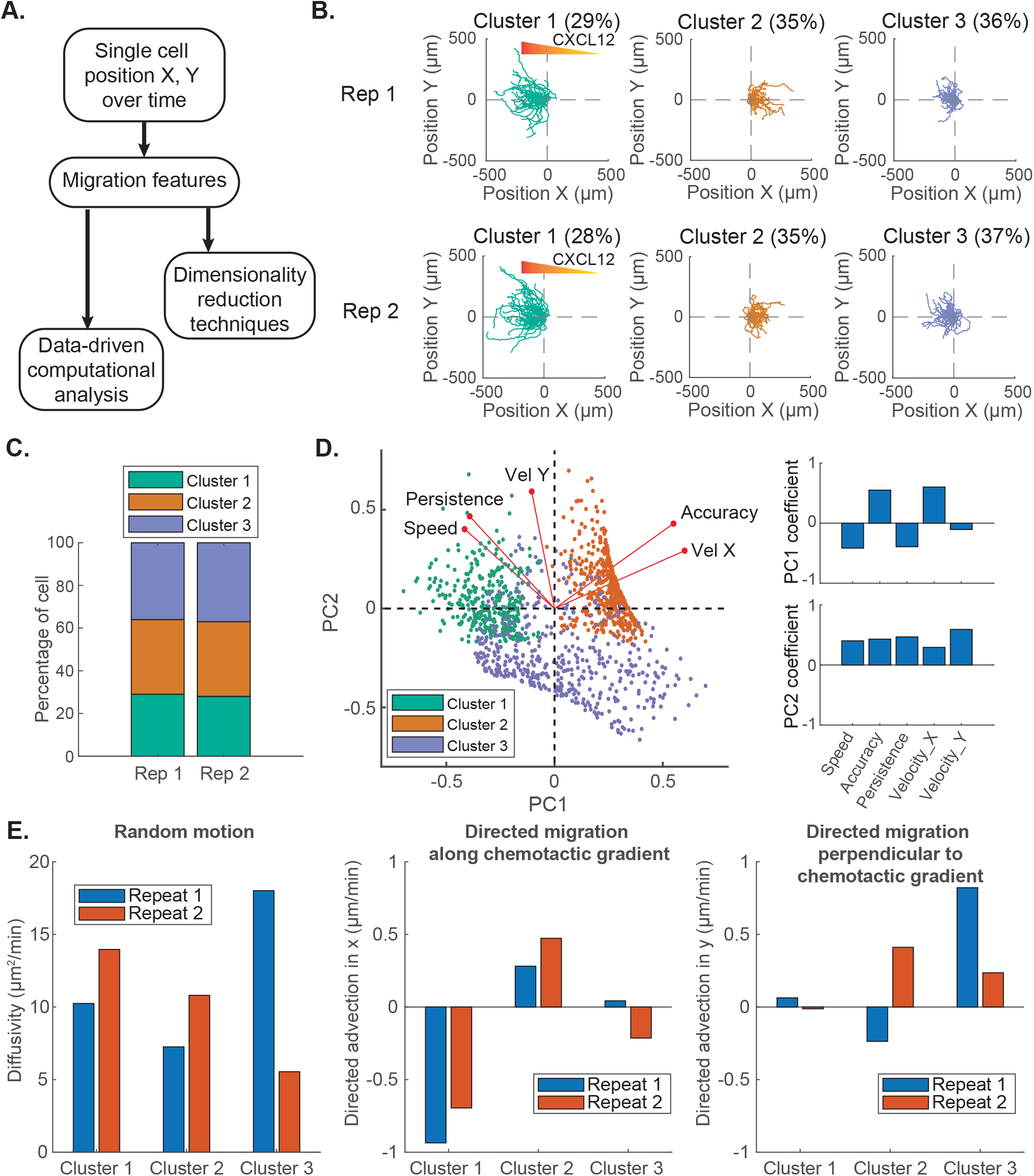
Separation and comparison of cell migration responses using migration features. **A**. Schematic showing the flow of quantitative analysis. Migration features are qualified from single cell positions over time and processed using dimensionality reduction techniques and data-driven computational analysis. **B**. Cells migrating in a CXCL12 gradient are clustered into three subpopulations using k-means clustering. 50 trajectories are randomly selected from each cluster. Trajectories are shown in different colors for each cluster. Clustering was conducted to two experimental repeats, shown in the first and second rows. **C**. Percentages of cells categorized to different clusters are similar in two repeats. **D**. Principal component analysis of five migration features: speed, persistence, accuracy, velocity in x and y. Right plots show first two principal component coefficients for each migration features. Left plot shows principal component scores for each cell (categorized by their cluster). The direction and length of the red vector indicate how each migration features contributes to the two principal components in the plot. **E**. Bar graphs showing the data-driven variational system inference of random motion, directed migration along and perpendicular to chemotactic gradient learned from different clusters of two replicates. They are represented by diffusivity, directed advection in x and y respectively. (Data includes two replicates, total of >1000 cells)

Accuracy and velocity in X direction are the main difference between correctly (cluster 1) and oppositely (cluster 2) migrating cells, while speed, persistence, and velocity in Y direction are unique in non-migrating cells (cluster 3) (**Fig. 2D**). Persistence describes whether cells are polarized or not, which relates to random motions of cells (28). Therefore, we speculate whether non-migrating cells have higher random motion comparing to migrating cells. Surprisingly, data-driven variational system inference reveals cells in all three clusters have similar random motion (characterized by diffusivity), while the main difference between clusters is directed migration along x direction (**Fig. 2E**). In summary, migrating cells exhibit faster and more persistent movement than non-migrating cells, but both show similar random motion.

### ERK/Akt amplitudes correlate with migration speed

Next we focus to identify unique signaling determinants for successful chemotaxis. Our previous work showed that absolute levels of ERK and Akt, known for driving cell migration and chemotaxis, are one of the descriptor of heterogeneity and regulated by pre-existing cell states (5). Naturally, we speculate whether heterogeneity of cancer cell migration is regulated by ERK and Akt signaling levels. Surprisingly, initial and average ERK or Akt activities do not correlate with migration responses (**Fig. 3A**) and have similar distributions among different clusters (**Fig. 3B**). Since we found that fluctuation amplitude, besides from absolute levels, is another descriptor of heterogeneity among individual cells, we hypothesize cells decode fluctuation amplitudes and frequencies to drive chemotaxis. Previous studies demonstrated ERK pulses with a range of pulsing frequencies (every few minutes to hours) regulating actin protrusions, collective wound healing, and drug resistance (12, 13, 16-18, 29-32). Therefore, we applied bandpass filtering to the signaling data to filter fluctuations at a specific range of frequencies (**Fig. 3C**). After exploring different ranges of frequencies, we found that the fluctuation amplitudes of the ERK or Akt signals filtered between 0.2-0.8 mHz correlate highly with the migration speed of the cell, where the correlation coefficients are 0.60 for ERK and 0.51 for Akt (**Fig. 3D-E**). Signal fluctuations between 0.2-0.8 mHz correspond to a fluctuation period of around 20-80 minutes, like the range of ERK fluctuations others have studied in migration (12, 16). While previous studies focused on either actin protrusions or average activities of ERK waves, our analysis discovered a correlation between signaling and migration outputs of each cell. Since migration speed is one of the dominant differences between migrating and non-migrating cells, we compared ERK and Akt fluctuation amplitudes between clusters. Migrating cells (cluster 1 and 2) have higher ERK and Akt amplitudes comparing to non-migrating cells (cluster 3) (**Fig. 3F**). Together, ERK and Akt fluctuation amplitudes, but not ERK and Akt absolute levels, are unique signaling determinants for successful chemotaxis.

**Figure 3:**
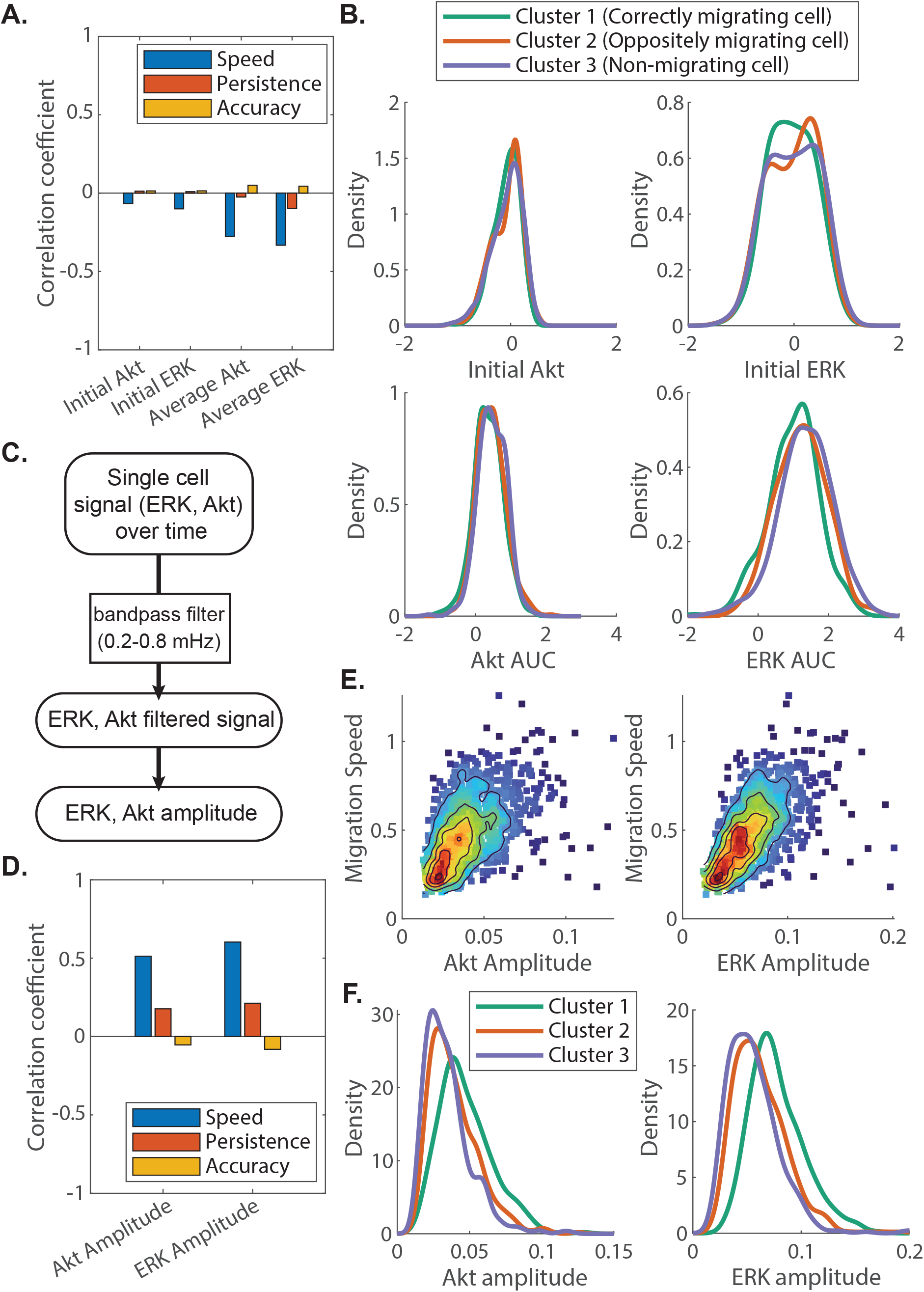
ERK and Akt signal fluctuation amplitudes, not absolute levels, correlate with migration responses. **A**. Correlation coefficients between ERK or Akt absolute signaling levels with migration features: speed, persistence, and accuracy. **B**. Kernel density estimation of probability density functions of initial Akt, initial ERK, normalized area under curve for Akt, and normalized area under curve for ERK for three different clusters. **C**. Schematic showing the signal processing method: single cell signal over time is passed through bandpass filter between 0.2-0.8 mHz and fluctuation amplitudes are calculated from filtered signal. **D**. Correlation coefficients between ERK or Akt amplitudes with migration features: speed, persistence, and accuracy. **E**. Density scatter plots of Akt and ERK amplitudes with migration speed show a linear relationship between Akt/ERK amplitudes and migration speed. **F**. Kernel density estimation of probability density functions of Akt and ERK amplitudes for three different clusters. (Data includes two replicates, total of >1000 cells)

### Lateral protrusion transition associates with chemotaxis

Cells facilitate migration towards chemoattractant through developing protrusions and retraction at distinct cell edges (33), which lead to dramatic changes in cell morphology. Since ERK and Akt signaling networks control cytoskeletal events resulting changes in cell morphology (12, 13), we ask if successful chemotaxing cells also have unique morphological changes. Previous study found that various cancer cell lines, including MDA-MB-231 breast cancer cell, in culture media exhibit sequential stages of movement which they termed ‘lateral protrusion’, ‘polarization’, and ‘tail retraction’ (34). They found that ‘lateral protrusion’ tends to occur with nucleus in the middle of two lateral protrusions and at 90° to the preceding tail retraction. Cells migrating with this sequential change in morphology (called ‘discontinuous mode’) are faster than ones without (called ‘continuous mode’). Using live-cell imaging of cell morphology, we observed MDA-MB-231 cells occasionally exhibiting lateral protrusion when they are migrating under CXCL12 gradient as if they are sensing the chemoattractant gradient before deciding the direction of movement (**Fig. 4A**). Therefore, we speculate to what extent heterogeneity of lateral protrusion transitions bridges ERK and Akt signal fluctuation and cancer cell migration.

**Figure 4:**
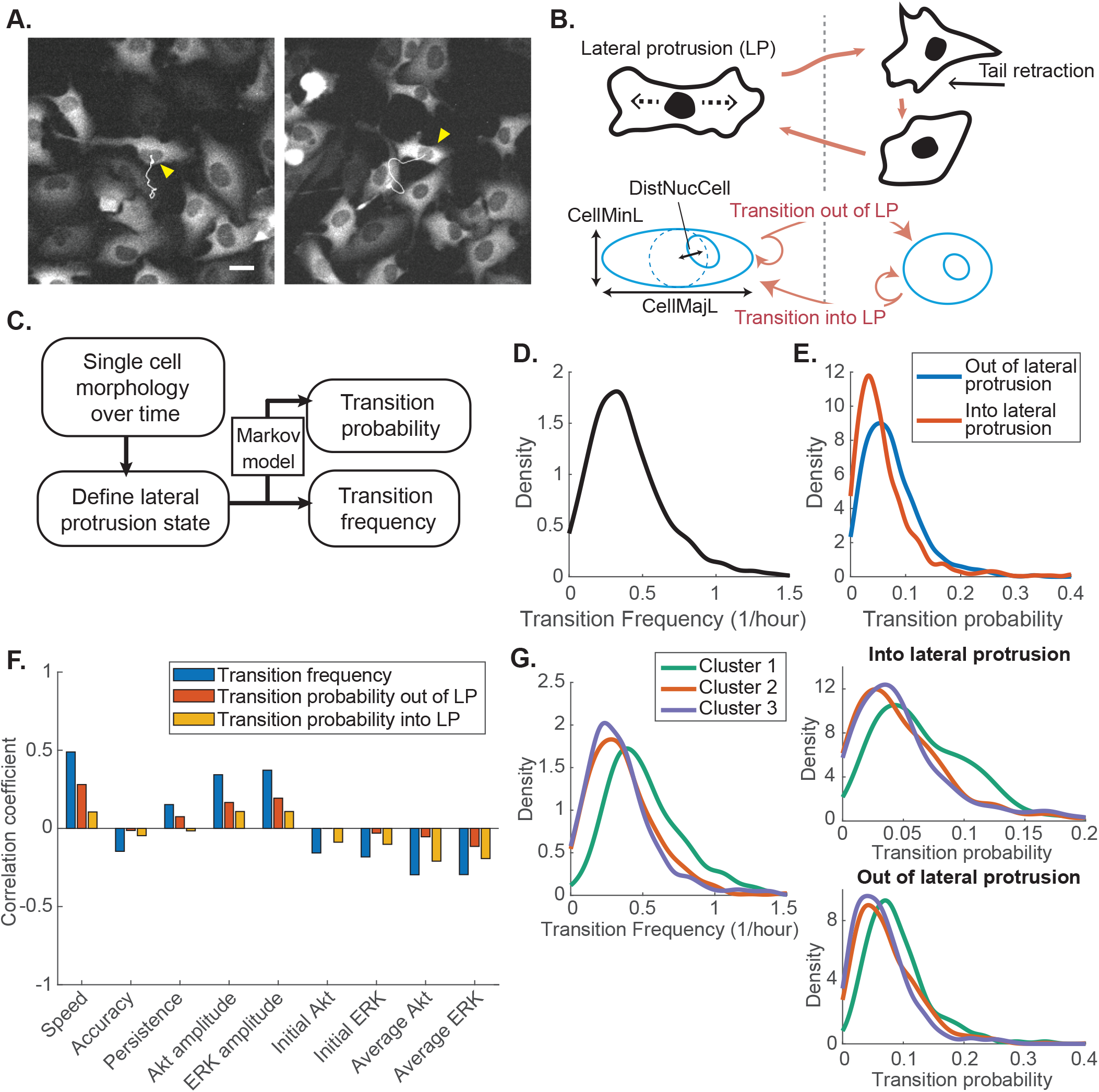
Lateral protrusion transition associates with chemotaxis responses. **A**. Two example pictures showing cell shape depicted by ERK KTR fluorescent signals. Yellow arrows point to two cells in the middle of each picture displaying lateral protrusion on both sides of the cell. Scale bar = 25µm. **B**. (Top) Schematic showing cells transitioning into lateral protrusion (LP) state and tail retraction state. Lateral protrusion is a general description of cell having two lateral protrusions with nucleus in the middle. (Bottom) Morphology features used to define lateral protrusion state. Cells can either stay at their current state or transition into or out of the lateral protrusion state. CellMajL is the cell major axis length, CellMinL is the cell minor axis length, and DistNucCell is the centroid distance between nucleus and cell. **C**. Schematic showing quantitative analysis method: single cell morphology features over time are passed through criteria to define lateral protrusion state. Transition frequency and transition probabilities were calculated using Markov model. **D-E**. Kernel density estimation of probability density functions of transition frequency (C) and transition probabilities (D) for the entire dataset. **F**. Correlation coefficients between migration or signaling features with transition frequency or transition probabilities. **G**. Kernel density estimation of probability density functions of transition frequency (left) and transition probabilities (right) for three different clusters. (Data includes two replicates, total of >1000 cells)

The lateral protrusion state is quantitatively defined by setting criteria with morphological properties such as aspect ratio, major and minor axis length, and positions of nucleus and cell (**Fig. 4B**). Since cells transition repeatedly into and out of lateral protrusion state, we used a Markov model for this behavior (**Fig. 4C**). Single-cell lateral protrusion transition frequency ranged from about once every hour to no transition within our observation time frame (**Fig. 4D**). The transition probabilities into and out of the lateral protrusion state also have a wide range of distributions in single-cell behaviors (**Fig. 4E**). The single-cell transition frequency and probability are not as correlated with migration/signaling features as between signaling fluctuation amplitudes and migration speed. We found that transition frequency correlates only fairly with ERK and Akt fluctuation amplitudes and migration speed, where the correlation coefficients are 0.37 for ERK, 0.34 for Akt, and 0.49 for speed (**Fig. 4F**). As a whole, the transition probabilities (red) do not correlate as well with ERK and Akt fluctuation amplitudes and migration speed, comparing to the transition frequency (blue). However, when we compare among clusters, interestingly, we found correctly migrating cells (cluster 1) have higher transition frequencies and probabilities compared to both oppositely migrating cells (cluster 2) and non-migrating cells (cluster 3) (**Fig. 4G**). This suggests that lateral protrusion transition associates with effective chemotaxis. This result is also in line with the possibility that cells are using the lateral protrusion state for sensing the CXCL12 gradient, as only the subpopulation of cells that correctly sensed and migrated towards CXCL12 has the highest frequency transitioning into lateral protrusion state.

### Temporal coupling among signaling, morphology, and migration

Cancer is known for having mutations and dysregulated signal networks, causing uncontrolled cell growth and migration. With ERK and Akt fluctuation amplitudes and morphological transition frequency correlating with migration speed, we speculate to what extent heterogeneity of network dynamics associates with migrating cell subpopulation and further investigate the dynamics and temporal coupling of ERK/Akt signaling, cell shape, and instantaneous migration speed. We first explored the dynamic relationship between ERK/Akt signals and cell aspect ratio of each cell. Cross-correlation analysis demonstrates migrating cells (cluster 1) have a lower correlation between ERK/Akt signal and cell aspect ratio and an increase in median time delay comparing to non-migrating cells (cluster 3) (**Fig. 5A-C**). This result indicates that migrating cells have a longer delayed and less controlled network coupling ERK/Akt signal with cell aspect ratio, comparing to non-migrating cells. Next, we also compared the dynamics of cell aspect ratio and instantaneous migration speed. In all three clusters, there are no median time delay between the aspect ratio and instantaneous speed. Cross-correlation analysis demonstrates migrating cells (cluster 1) have a small increase in correlation between cell aspect ratio and instantaneous speed, comparing to non-migrating cells (cluster 3) (**Fig. 5D-E**). This result indicates that migrating cells have a slight increase in regulation controlling cell morphology and migration speed. Together, these data suggest that although correctly migrating cells possess higher signaling fluctuation in ERK and Akt and higher morphological transition frequency, they have less correlated network coupling ERK/Akt signaling and cell aspect ratio, comparing to non-migrating cells. This implicates that processes connecting ERK/Akt signaling and morphological changes could potentially be tuned to control dysregulated cancer migration.

**Figure 5:**
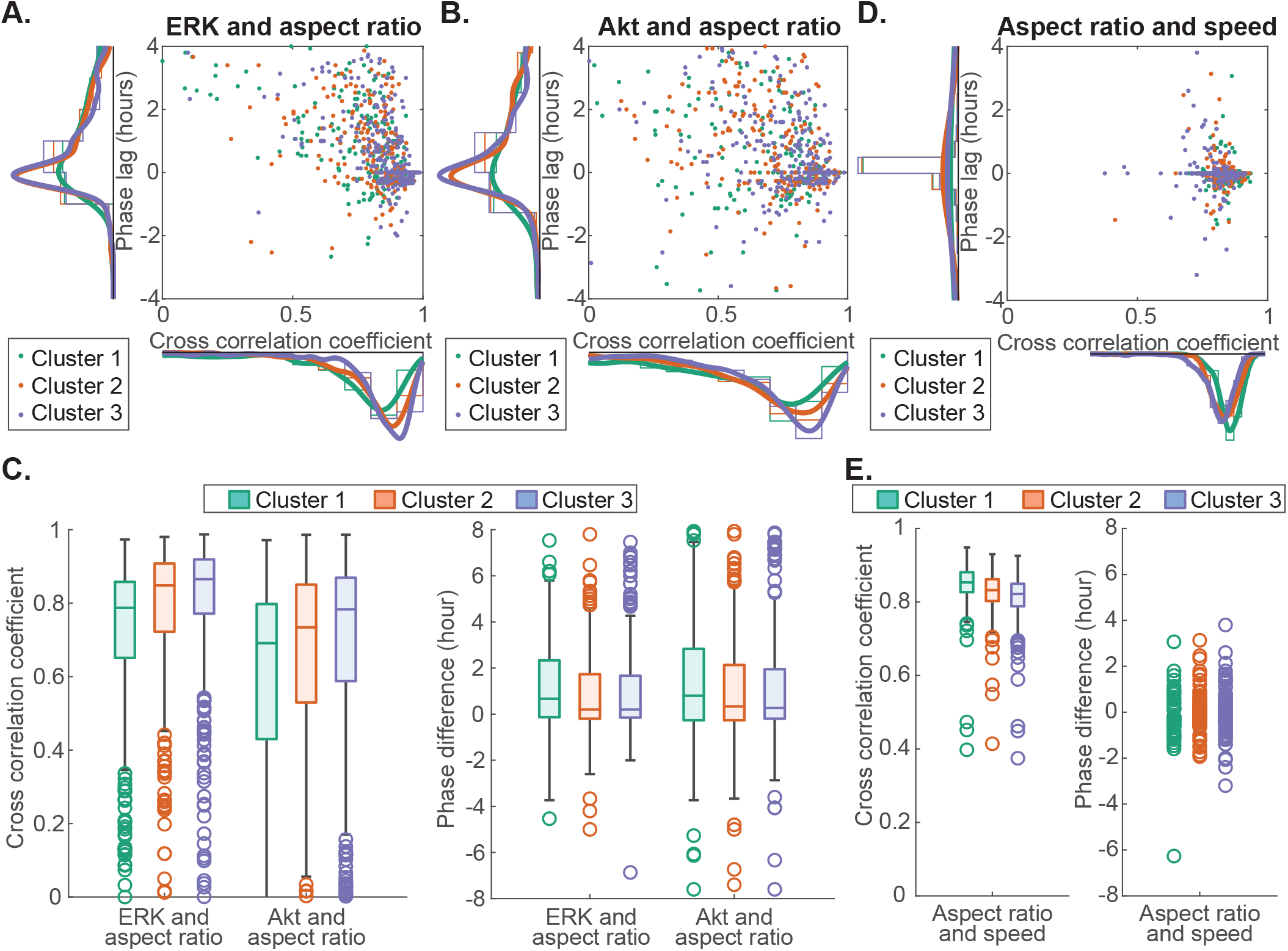
Temporal coupling among signaling, morphology, and migration. The ERK/Akt signaling, aspect ratio, and instantaneous speed of each cell are passed through cross-correlation analysis. Highest cross correlation coefficient and corresponding phase difference are captured. **A**. Kernel estimated probability density functions plotted with scatter plot of cross correlation coefficient and phase difference between ERK signaling and aspect ratio for different clusters. **B**. Kernel estimated probability density functions plotted with scatter plot of cross correlation coefficient and phase difference between Akt signaling and aspect ratio for different clusters. **C**. Box plot of cross correlation coefficient and phase difference between ERK/Akt signaling and aspect ratio to show the difference among clusters. **D**. Kernel estimated probability density functions plotted with scatter plot of cross correlation coefficient and phase difference between aspect ratio and instantaneous speed for different clusters. **E**. Box plot of cross correlation coefficient and phase difference between aspect ratio and instantaneous speed to show the difference among clusters. (Data includes two replicates, total of >1000 cells)

## Discussion

Resolving dynamic and integrated processes in chemotaxis at single-cell resolution offer insights in understanding heterogeneity in cancer cell migration and metastasis. In this study, we combined quantitative analysis along with live-cell imaging and automated image analysis to collect an integrated dataset and extract biological understanding associated with cell-to-cell variability in signaling, morphology, and chemotaxis responses. Through a combination of time-dependent data processing, dimensionality reduction techniques, and data-driven inference, we discovered unique properties in migratory cells comparing to non-migratory cells. Migrating cells have higher ERK/Akt fluctuation amplitudes, more frequent morphological transition, but less correlated network coupling ERK/Akt signaling and morphological changes. To our knowledge, this is the first report demonstrating dynamic network coupling signaling and morphology as key determinants of successful chemotaxis.

Our group previously demonstrated that activations of ERK and Akt differ among cells in response to CXCR4 signaling and other signaling molecules (5, 7). The signaling heterogeneity is regulated by pre-existing cell states, determined by Ras, mTORC1, and PI3K activities. However, the heterogeneity of processes connecting signaling and chemotaxis remain unknown. Previous studies showed that PI3K and MAPK signaling are connected with feedbacks to reorganization of actin cytoskeleton and resultant changes in cell morphology, which is known to regulate cell migration. Due to the network feedbacks, ERK pulses have been showed to regulate protrusion events (12), collective migration (16, 18), oncogenic transformation (13), and drug resistance (17). These studies are consistent with our findings but we first report that ERK and Akt fluctuation amplitudes, rather than absolute levels of ERK/Akt signaling, correlate with heterogeneous migration speed. We further demonstrate that ERK/Akt fluctuation amplitudes are key factors for effective chemotaxis, building a connection between signaling and chemotaxis heterogeneity. Another study demonstrated that cells migrating with consistent change in cell morphology have higher speed than ones with more uniform cell shape (34). They characterized cells with consistent change in cell morphology transitioning into lateral protrusion state. This study is consistent with our finding but we first report that transition frequency into and out of lateral protrusion state correlates with heterogeneous migration speed. We further demonstrate that morphology transition is important for cells to correctly sense and migrate towards CXCL12, establishing morphological transition as a potential bridging point between signaling and chemotaxis heterogeneity.

Ras/PI3K/ERK signaling network has been studied extensively to dissect the general signaling network coupling protrusion activities (12, 13, 29-32). This network couples with cytoskeletal signals and is triggered by localized protrusion activities with ERK pulses lagging protrusion events (12). In our finding, we found that ERK signaling cross correlate with cell shape with a wide range of phase difference. While a group of cells demonstrate ERK signal lagging morphological change, our single cell analysis also reported another group of cells with ERK signal leading morphological change. This demonstrates a population of cells may have heterogeneous network coupling signaling and morphology. We also reported the network coupling ERK/Akt signaling and morphological changes is less correlated for migrating cells comparing to non-migrating cells. We believe that feedback networking wiring Ras/PI3K/ERK and morphological changes in migrating cells could be altered and have a different feedback delay in the Ras/PI3K/ERK signaling network from non-migrating cells. Previous evidence showed that B-Raf decay rate in Ras-ERK pathway reshapes downstream transcription and cell fates (35), suggesting possible difference in kinase deactivation rate contributing to heterogeneous network connection and migration response. Further, we extended our work to compare the dynamics of morphological change and the resulting instantaneous migration speed. We found that cell shape and instantaneous speed are well cross correlated with no phase difference. Since our study compared the difference in a thousand of cells, we show that good migrators have small increase in cross-correlation than non-migrating cells. To our knowledge, this is the first report that cells have different instantaneous speeds when they change their cell shape. With lateral protrusion being one of the morphological transition during chemotaxis, we propose that cells might use lateral protrusion to help sense chemotactic gradient and reorient their direction of motion. As a result, the better the cell can use this mechanism, the better they are in migrating.

In this study, the characterization of network dynamics is limited by the imaging frequency and the types of live-cell reporters in the cell. Current imaging frequency is 5 times faster than the signal dynamics that we studied. With recent advances in automated imaging techniques and live-cell reporters, the imaging frequency and number of live-cell reporters can be increased in the future without causing phototoxicity or fluorescence crosstalk. This improvement will help reveal other mechanisms regulating heterogeneity in cancer cells. Besides, cells migrate differently in 2D vs 3D environments. We will need to confirm if the discoveries that we found in this 2D chemotaxis study are applicable in a 3D environment. Our quantitative analysis approach in this study was focused to find relationship among signaling, morphology, and migration and discover variations between migrating and non-migrating cells. While we found interesting interconnections in signaling, cell shape, and movement in migrating cells, future studies are needed to identify molecular mechanisms regulating these connections. Our data also suggest future explorations of pre-existing cell states with unique feedback delay in Ras/PI3K/ERK signaling network regulating migration heterogeneity.

Our study design combines live-cell imaging and automated image analysis to capture large scale, multidimensional data. With the integrated quantitative analysis used in this study, we revealed dynamic network coupling signaling and morphology as key factor for effective chemotaxis. Our unique approach paves the path towards better understanding of mechanisms regulating heterogeneous responsiveness in cancer cells and future identification of new interventions controlling cancer metastasis.

## Method

### Cell culture

We purchased breast cancer cell lines MDA-MB-231 cells from ATCC (Manassas, VA) and cultured them in Dulbecco’s Modified Eagle Medium (DMEM) supplemented with 10% fetal bovine serum (FBS), 1% penicillin/streptomycin (Pen/Strep) (Thermo Fisher Scientific, Waltham, MA), 1% GlutaMAX (Thermo Fisher Scientific). We maintained cells at 37 °C in a humidified incubator with 5% CO_2_. Prior to loading cells for chemotaxis experiment, we precultured MDA-MB-231 cells in FluoroBrite Full glucose imaging media, 10% FBS, 1% Pen/Strep, and 1% GlutaMAX for 2 days.

### Stable expression of fluorescence reporters in cells

We used a previously constructed PiggyBac transposon vector (pHAEP) containing histone 2B fused to mCherry (H2B-mCherry), Akt-KTR (Aquamarine), ERK-KTR (mCitrine), and a puromycin selection marker all separated by P2A linker sequences (5). We cotransfected the pHAEP transposon and Super PiggyBac transposase vector (System Biosciences, Palo Alto, CA, USA) into MDA-MB-231 breast cancer cells. We selected stable cells with puromycin (4µg/ml). We verified expression of the full pHAEP reporter construct by fluorescence microscopy. Additionally, the cells are overexpressed with receptor CXCR4 so that they will migrate towards the chemoattractant. We transduced MDA-MB-231 cells stably expressing the pHAEP reporter with lentiviral vector for CXCR4.

### Chemotaxis experimental setup

We used ibidi chemotaxis microfluidic device (µ-Slide chemotaxis, ibidi, Planegg. Germany) with two large reservoirs containing different concentrations of CXCL12 at both sides of the chambers. We cultured MDA-MB-231 cells in the chemotaxis device under FluoroBrite Full glucose media (3% FBS, 1% Pen/Strep, and 1% GlutaMAX) for two hours and introduced a chemoattractant gradient with ligands CXCL12 (300ng/ml to 0ng/ml) (from R&D Systems, Minneapolis, MN). We took time-lapse multi-color fluorescence images of the chemotactic cells using a fully automated EVOS M7000 imaging system (ThermoFisher) at 4-minute time intervals for 24 hours. The imaging system is equipped with onstage incubator and autofocusing for live-cell imaging.

### Automated time-lapse fluorescence imaging and image processing

After taking time-lapse multi-color fluorescence images, images were processed automatically using our custom MATLAB code. We made advancement from our program previously described (5). Briefly, our program segments nuclei and cell cytoplasms using adaptive thresholding to detect the sharp increase in fluorescence intensity at the edges. It then conducts a segmented nuclei assisted watershedding to segment merged cell cytoplasm. Further, it extracts position and morphology properties of the nucleus and cell and quantify the ERK and Akt KTR values as log2 of the cytoplasmic to nuclear KTR fluorescence intensities. Cells are connected between time points using segmented nuclear position. Our program identifies high-confident cell matches and other scenario such as mitosis, missed frame, and merged mask using statistical approach based on nuclei distance and morphological properties. The tracks are trimmed when the cells are dividing because the signaling and morphology data is not accurate during mitosis.

### Separation and comparison of cell’s migratory behaviors: clustering and principal components analysis

We used unsupervised k-means clustering algorithm to separate single-cell tracks into three clusters by minimizing within-cluster variances in all five migration features: speed, persistence, accuracy, and velocity in x and y directions. We chose three clusters because that best separate cells based on their trajectories. We combined single-cell tracks from two experimental replicates and normalized every migration feature. Then, we used MATLAB to conduct k-means clustering. Cluster ID of individual cell was used throughout the entire analysis.

We compared the contribution of five migration features in separating the three clusters by performing a weighted principal components analysis. We used inverse variances of the migration features as weights. We visualized the orthonormal principal component coefficients for each migration feature and principal component scores for individual cell in a single plot. Cluster ID of each cell is also indicated by different color. The direction and length of each migration feature vector indicate how each feature contributes to the separation of the three clusters.

### Comparison of cell’s migratory behaviors: variational system inference

We used previously developed method, variational system inference, to identify the random and directed motion of cells in each cluster. Briefly, with large enough length scales, we studied individual cells as densities or continuum fields and characterized the motion of cells as change in densities. We used an advection-diffusion partial differential equation to model cells in continuum fields. The diffusive system at large length scale is the behavior of cells taking random walk. Advection represents the homogenized behavior of cells with a directed motion. A higher advective behavior in comparison to diffusive means a more persistent behavior in cells. We treated the densities of cells in each cluster independently to determine their behavior. The parameters are estimated using the variational system inference approach. Details of this method are presented in our previous work with brief details in the supplementary information (36-38).

### Quantification of signaling fluctuation amplitudes: bandpass filtering

We quantified ERK and Akt signaling fluctuations between 0.2 to 0.8 mHz using bandpass filtering. We specified the filtering frequencies and imaging frequency and applied bandpass filtering to ERK or Akt signals using MATLAB function. Then, we identified the local maxima and minima of the filtered signal and computed the average amplitudes.

### Quantification of morphological transitions

We defined lateral protrusion state using morphological properties such as aspect ratio, major and minor axis length, and positions of nucleus and cell. Lateral protrusion state has an aspect ratio larger than 2 and a centroid distance between nucleus and cell within half of minor axis length. We used Markov model to describe this morphology transition behavior. A Markov model assumes that future states depend only on the current state, not on any past events. We counted the number of times cells transition into and out of lateral protrusion state, divided it by length of cell track, and converted into transition frequency for each cell. We also calculate the average transition probabilities into or out of lateral protrusion state based on the number of time cells stayed in the same state vs number of time cells changed state.

### Temporal coupling among signaling, morphology, and migration

We analyzed the temporal coupling using cross-correlation analysis. Cross-correlation analysis was conducted on raw ERK and Akt signals, aspect ratio representing cell morphology change, and calculated instantaneous speed based on cell nuclear positions over time. We first calculated the phase differences and cross correlation coefficient of each cell, then identified the maximum cross correlation coefficient and the corresponding phase difference. The maximum cross correlation coefficients and corresponding phase differences of each cell were compared among different clusters to find any difference in temporal coupling for migrating cells.

## Acknowledgements

This study was supported by NIH grants R01CA238042, U01CA210152, R01CA238023, R33CA225549, R50CA221807, and R37CA222563. Research also was supported by funding from the W.M. Keck Foundation.

## Author Contributions

K.E.L and G.D.L. are senior coauthors. K.K.Y.H., P.C.K., G.D.L., and K.E.L. participated in research design. K.K.Y.H. conducted experiments. K.K.Y.H., S.S., and K.G. performed the data analysis. K.K.Y.H., S.S., G.D.L. and K.E.L. wrote the manuscript. All authors contributed to editing the article and approved the submitted version.

## Figure Legends

**Figure S1:**
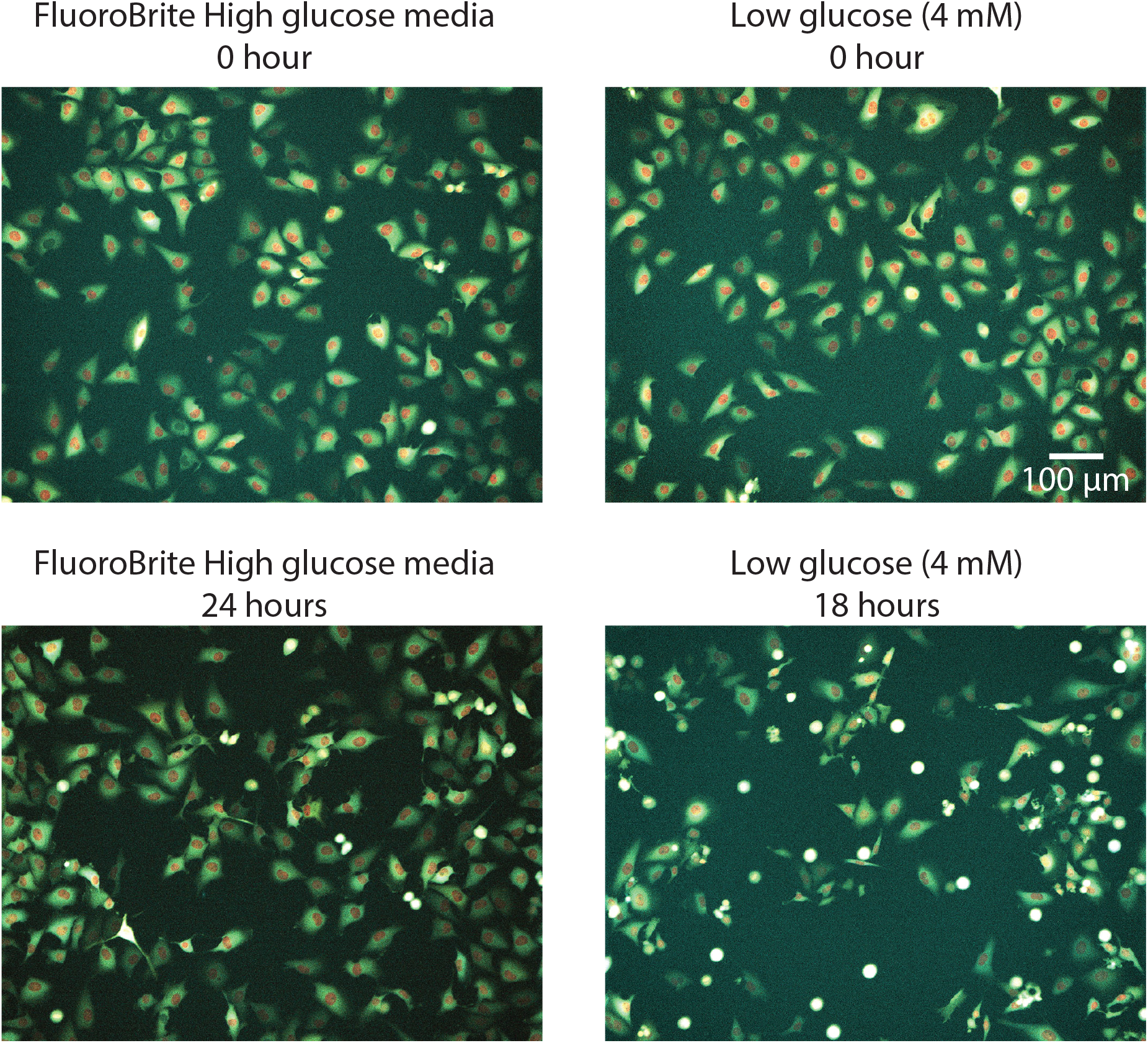
Long term cell migration in microfluidic device under regular fluorescence imaging. Sample images with combined colors of H2B-mCherry (Red), Akt-KTRs (Cyan), and ERK-KTRs (Yellow). Cells are healthy in the microfluidic device with high glucose before (Top left) and after 24 hours of imaging at 4-minute time interval (Bottom left). When low glucose media was used, cells started dying inside the microfluidic device after 18 hours of fluorescence imaging (Right).

