## Supplementary information for "Cell-to-cell variability of dynamic CXCL12-CXCR4 signaling and morphological processes in chemotaxis"

### 1 Variational System Inference

The general strong form for the Advection-Diffusion PDE is

$$\frac{\partial C}{\partial t} = D \cdot \Delta C - \cdot \nabla \cdot C[v_x, v_y] \tag{1}$$

where $D$, $v_x$ and $v_y$ are the Diffusivity and the Advective velocities in x and the y directions, respectively.

$$\begin{aligned}\int_\Omega \frac{\partial C}{\partial t} w d\Omega &= \int_\Omega \left(\nabla \cdot (D\nabla C) - \nabla \cdot (C\boldsymbol{v})\right) w d\Omega \\ &= \int_\Omega \left(-D\nabla w \cdot \nabla C + C\boldsymbol{v} \cdot \nabla w\right) w d\Omega + \int_{\partial\Omega} w \left(D\nabla C - C\boldsymbol{v}\right).\boldsymbol{n} ds \end{aligned} \tag{2}$$

where $\boldsymbol{w}$ represents the weighting function (an arbitrary variation on the density field). Following the standard procedure in finite element analysis (FEA) computations of discretizing the domain and accounting for the arbitrariness of the weighting function's degrees of freedom, Equation (2) leads to a set of algebraic equations written as the residual vector, $\boldsymbol{R} = \boldsymbol{0}$ where the residue is given as :

$$\boldsymbol{R} = -\boldsymbol{A}_e \left[\int_{\Omega_e} \left(\frac{\partial C}{\partial t}\boldsymbol{N} + D\nabla\boldsymbol{N} \cdot \nabla C - C\boldsymbol{v} \cdot \nabla\boldsymbol{N}\right) d\Omega - \int_{\partial\Omega_e} \boldsymbol{N} \left(D\nabla C - C\boldsymbol{v}\right).\boldsymbol{n} ds\right] \tag{3}$$

where $\boldsymbol{A}_e$ is the FEM assembly operation and $\boldsymbol{N} = [N^1, \cdots, N^n_{nel}]$ is the vector of interpolation functions in an element. The residue can be succinctly written as:

$$\boldsymbol{R}(D, v_x, v_y) = y - \begin{bmatrix}\Xi_1, \Xi_2, \Xi_3\end{bmatrix} \begin{bmatrix} D \\ v_x \\ v_y \end{bmatrix}$$

where

$$y = -\boldsymbol{A}_e \int_{\Omega_e} \frac{\partial C}{\partial t} \boldsymbol{N} d\Omega$$

$$\Xi_1 = \boldsymbol{A}_e \left[ \int_{\Omega_e} \nabla \boldsymbol{N} \cdot \nabla C d\Omega - \int_{\partial\Omega_e} \boldsymbol{N} \nabla C . \boldsymbol{n} ds \right]$$

$$\Xi_2 = \boldsymbol{A}_e \left[ \int_{\Omega_e} -C \frac{\partial N}{\partial x} d\Omega + \int_{\partial\Omega_e} N C n_x ds \right]$$

$$\Xi_3 = \boldsymbol{A}_e \left[ \int_{\Omega_e} -C \frac{\partial N}{\partial y} d\Omega + \int_{\partial\Omega_e} N C n_y ds \right]$$

The Variational System Identification problem is then posed as the following minimization problem:

$$D, v_x, v_y = \underset{\widetilde{D}, \widetilde{v}_x, \widetilde{v}_y}{\arg\min} |\boldsymbol{R}(\widetilde{D}, \widetilde{v}_x, \widetilde{v}_y)|^2 \quad (4)$$
